# Genetic variability, including gene duplication and deletion, in early sequences from the 2022 European monkeypox outbreak

**DOI:** 10.1101/2022.07.23.501239

**Authors:** Terry C. Jones, Julia Schneider, Barbara Mühlemann, Talitha Veith, Jörn Beheim-Schwarzbach, Julia Tesch, Marie Luisa Schmidt, Felix Walper, Tobias Bleicker, Caroline Isner, Frieder Pfäfflin, Ricardo Niklas Werner, Victor M. Corman, Christian Drosten

**Affiliations:** Institute of Virology, Charité – Universitätsmedizin Berlin, corporate member of Freie Universität Berlin, Humboldt-Universität zu Berlin, and Berlin Institute of Health, 10117 Berlin, Germany; German Centre for Infection Research (DZIF), partner site Charité, 10117 Berlin, Germany; Centre for Pathogen Evolution, Department of Zoology, University of Cambridge, Downing St., Cambridge, CB2 3EJ, U.K.; Klinik für Innere Medizin - Infektiologie, Vivantes Auguste-Viktoria-Klinikum, Rubensstr. 125, 12157, Berlin, Deutschland; Department of Infectious Diseases and Respiratory Medicine, Charité – Universitätsmedizin Berlin, corporate member of Freie Universität Berlin, Humboldt-Universität zu Berlin, and Berlin Institute of Health, 13353, Berlin, Germany; Division of Evidence-Based Medicine (dEBM), Department of Dermatology, Venereology and Allergy, Charité-Universitätsmedizin Berlin, Corporate Member of Freie Universität Berlin, Humboldt-Universität zu Berlin, and Berlin Institute of Health, Berlin, Germany; Labor Berlin – Charité Vivantes GmbH, 13353 Berlin, Germany

## Abstract

Genome sequences from 47 monkeypox virus infections detected in a German university virology laboratory were analyzed in context of other sequences from the 2022 outbreak and earlier monkeypox genomes. Identical non-synonymous amino acid changes in six genes and the signature of APOBEC editing match other sequences from the European outbreak. Non-synonymous changes that were present in one to three sequences were found in 34 other genes. In sequences from two lesions of one patient, an 856 nucleotide translocation between genome termini resulted in the duplication of an initial 5’ gene, and the disruption or complete deletion of four genes near the 3’ genome end. Orthopoxvirus genome rearrangements of this nature are known to confer fitness advantages in the face of selection pressure. This change may therefore represent an early virus adaptation in the novel widespread and sustained human-to-human context of the current monkeypox outbreak.

## Introduction

Monkeypoxvirus (MPXV) is a species of the Orthopoxvirus genus in the *Poxviridae* family of large double-stranded DNA viruses. Prior to the 2022 outbreak, the virus was endemic in at least 10 central and western African countries (DRC, CAR, Nigeria, Republic of the Congo, Sierra Leone, Cameroon, Côte d’Ivoire, Gabon, Liberia, South Sudan)^1^. Phylogenetic analyses show central and western African MPXV clades^2^. Western clade cases are rarely fatal in non-immunocompromised patients, while the central African strain is more virulent, with case fatality rates typically in the range 1% to 5% although occasionally higher^3^. The number of reported confirmed and probable human cases have increased every decade since the first human case was detected in 1970 (1970s: 47, 1980s: 356, 1990s: 520, 2000s: 10,166 (including suspected cases), 2010s: 19,068 (including suspected cases)^1^. Over 99% of all cases have been attributed to the central African clade^1^. Cases are ascribed to zoonoses, with sleeping on the ground and living close to or visiting the forest being reported as risk factors for acquiring MPXV infection^1^. The number and identity of MPXV reservoir species is unknown, despite considerable effort^4–10^. Human transmission chains have increased in length over the decades, but until recently their maximum lengths have been estimated at approximately seven^3,11^.

Variability in orthopoxvirus gene composition is well established, observed in both short-term time spans such as cell culture passage and short-lived human MPXV outbreaks^12–15^ and also in the general long-term pattern of gene inactivation and loss during orthopoxvirus evolution^16,17^. Genome plasticity is especially pronounced in two inverted terminal repeat (ITR) regions at the genome termini^18,19^. ITR regions contain predominantly immunomodulatory host range factors that also impact virulence^16,20–22^. Modification of these regions is considered a primary mechanism of rapid orthopoxvirus adaptation following host switches^12,13^, including in human MPXV cases in central Africa^20,23^. Deletions and insertions in the terminal genome regions are prominent differences between the western and central African MPXV clade sequences^24^ as also found in sequences from the US outbreak in 2003^2^. Recombination-mediated genome expansion and contraction through gene duplication, with subsequent gene deletion following acquisition of beneficial mutations, has been convincingly suggested as an “accordion”-like mechanism of rapid poxvirus adaptation following host switches^12,13^. The expansion increases the genome surface area available for beneficial mutations, temporarily compensating, in specific genes, for the low mutation rate and overall genetic stability of the rest of the double-stranded DNA genome, reminiscent of the genetic flexibility in bacterial evolution^12^. The pattern of rapid genetic change following host switching is thought to result from the opportunity for the virus to specialize in a new host, wherein specific host range genes required in the previous host(s) can be discarded or repurposed. The long-term pattern of gene inactivation and loss observed in orthopoxviruses is likely a result of host specialization.

In May 2022, new cases of MPXV were reported in the UK^25^. By July 14, 2022, cumulative confirmed cases had risen to 10,845 across 59 countries that had not historically reported cases^26^. The widespread MPXV outbreak, with its unprecedented human-to-human transmission, holds the potential for rapid adaptation to the human host. Any such changes in 2022 outbreak sequences should therefore be closely monitored and assessed for possible phenotypic effects. Here we present genetic analysis of 47 cases from Berlin, Germany, sampled between May 20 and July 4, 2022, including gene duplication and deletion in the sequence of one patient.

## Methods

We used swabs taken from lesions and a dual-target PCR strategy, with two independent gene regions detected subsequently in two separate reactions or simultaneously in one tube. This strategy involved one assay for generic testing for Orthopoxvirus^27^ and a MPXV-specific assay^28^. Assays were commercially available kits (LightMix^®^ Modular Virus kits, TIB Molbiol, Berlin; Germany), including internal amplification controls to test for PCR inhibition. Total nucleic acids (NA) were extracted using the Roche MagNAPure 96 and the Viral NA Small Volume Kit (Roche, Mannheim, Germany) or QIAamp Viral RNA Kits (Qiagen, Hilden, Germany). For high-throughput sequencing, library preparation was performed using the KAPA DNA Hyper Prep Kit (Roche Molecular Diagnostics, Basel, Switzerland) according to manufacturer’s instructions. DNA libraries were sequenced on Illumina MiniSeq (300 cycles, paired-end) or Illumina NextSeq (150 or 300 cycles, paired-end). Sequencing reads were aligned to the full-genome outbreak sequence ON563414.2 using Geneious (version 2022.0.1), which was then used to call consensus sequences.

We downloaded full-genome MPXV sequences from the US National Center for Biotechnology Information (NCBI) on June 14, 2022 using the query ‘monkeypox[All Fields] AND “Monkeypox virus”[porgn] AND “complete genome”[All Fields]’. After visual examination of an alignment of all sequences, seven sequences (HQ857563.1, KC257459.1, KJ642612.1, KJ642614.1, KJ642618.1, MN346693.1, and MT903341.1) were omitted from further analysis due to their degree of divergence from all other full-genome MPXV sequences and from each other. This resulted in a set of 88 full genome pre-2022-outbreak sequences. We also downloaded^29,30^ 83 sequences from 14 countries associated with the 2022 outbreak (with counts): Australia (1), Belgium (1), Finland (2), France (1), Germany (25), Israel (1), Italy (3), Netherlands (1), Portugal (28), Slovenia (1), Spain (4), Switzerland (2), United Kingdom (5), USA (8). For accession numbers, see Table S1.

We examined MPXV patient genome sequences for single-nucleotide polymorphisms and performed a gene-level analysis, looking for non-synonymous substitutions, changes in start/stop codons, and insertions or deletions. The conserved central region of the MPXV genome (locations 56,000 to 120,000)^20^ was used to infer a maximum likelihood phylogenetic tree using iqtree2 (version 2.2.0)^31^ with a K3Pu+F+I model (determined via ModelFinder^32^) and ultrafast bootstrapping^33^ (1000 bootstrap trees), based on an alignment constructed using mafft (version 7.487)^34^.

## Results

We confirmed MPXV infection in swab samples sent in from 47 male patients aged 23-58. For all samples, specific MPX testing was requested due to medical history suggestive of MPXV infection, including lesions (mainly) in the perianal/urogenital regions, genital rash, fever, and malaise. Swab samples from lesions were taken between May 20 and June 28, 2022 in outpatient departments in Berlin. All 47 samples were RT-PCR positive according to both the generic orthopoxvirus assay and the MPXV-specific assay. High-throughput sequencing resulted in genome coverage of 99.95% or higher for 38 samples, and 95.0 to 99.5% coverage for the remaining ten (two lesions from one of the patients were sequenced, as discussed below) (Table S2).

A phylogenetic tree containing pre-outbreak sequences (n=88), 2022 outbreak sequences from others (n=83), and our 48 sequences (219 in total) resulted in a grouping of the European outbreak sequences in a monophyletic clade (Fig. 1). The nearest pre-2022-outbreak sequence is a human sequence (ON676708.1) detected in the US in a traveler from Nigeria, collected in 2021^35^. The sequences from Berlin are widely spread in clade 3, consistent with multiple introductions and a high diversity of MPXV in the city.

**Figure 1:**
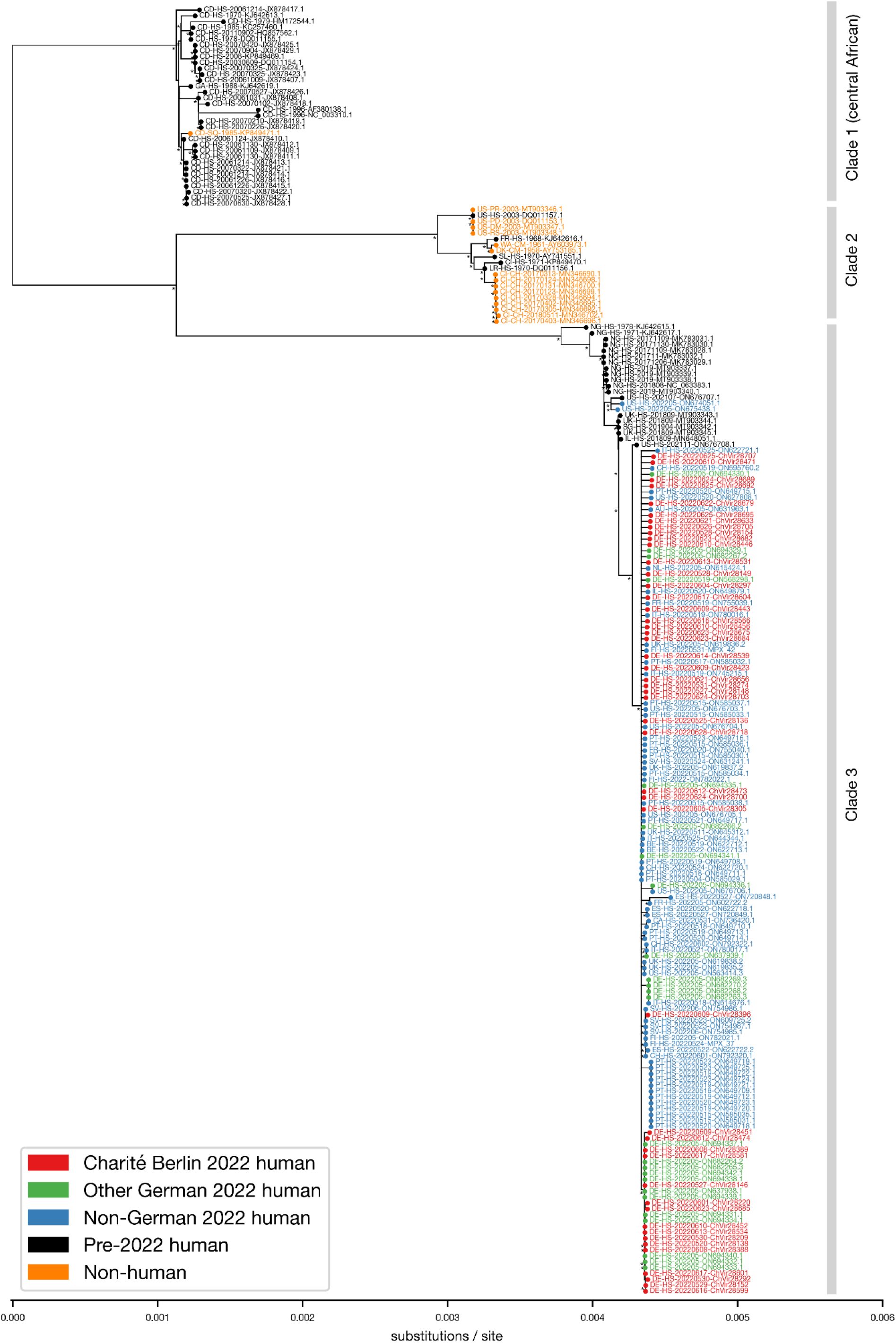
Maximum likelihood tree. Outbreak sequences (n=131) and all pre-outbreak NCBI full genome MPXV sequences (n=88). Tip names are composed of four hyphen-separated fields: a two-letter country or region code, a two-letter host species code, a YYYYMMDD date (or some fraction thereof when only partially known), and an identifier. The identifier is either a GenBank accession number and version or a sequence from this paper starting with the prefix “ChVir” in red text. Country/region abbreviations (count): AU: Australia (1), BE: Belgium (2), CA: Canada (1), CD: Democratic Republic of the Congo (33), CH: Switzerland (4), CI: Cote d’Ivoire (10), DE: Germany (73), DK: Denmark (1), ES: Spain (4), FI: Finland (4), FR: France (4), GA: Gabon (1), IL: Israel (2), IT: Italy (6), LR: Liberia (1), NG: Nigeria (12), NL: Netherlands (1), PT: Portugal (28), SG: Singapore (1), SL: Sierra Leone (1), SV: Slovenia (5), UK: United Kingdom (8), US: USA (15), WA: West African (1). Host abbreviations (count): CH: pan troglodytes verus (9), CM: cynomolgus monkey (2), DM: dormouse (1), HS: homo sapiens (203), PD: prairie dog (1), PR: cricetomys gambianus (1), RS: rope squirrel (1), SQ: squirrel (1). Tree made with iqtree2^31^ with the K3Pu+F+I model (as selected by ModelFinder^32^) and ultrafast bootstrapping^33^ (1,000 bootstrap trees). Branches with bootstrap support < 30 are collapsed and those with bootstrap support >= 70 are indicated by an asterisk.

Our 48 outbreak sequences were individually compared to the closest reference and also to each other. Nucleotide sequence identity was above 99.85% in all comparisons (Digital Supplement), with the exception of sequence ChVir28389, discussed below. In agreement with the observation of others^36,37^, the pattern of nucleotide (nt) changes in the Berlin outbreak sequences shows a strong bias towards GA→AA and (the complementarily equivalent) TC→TT substitutions.

### Non-synonymous gene substitutions

We examined the 38 highest coverage (>99.95%) genomes generated here for unambiguous evidence of non-synonymous substitutions. All 38 sequences have identical non-synonymous changes in six genes: E353K in MPXVgp045; D88N in MPXVgp078; M142I in MPXVgp079; E162K in MPXVgp083; H221Y in MPXVgp157; and P722S in MPXVgp182. Gene names are from the GenBank annotation of the closest pre-outbreak genome, ON676708.1, with notes and frequently-used alternate names, usually of homologous genes in vaccinia virus Copenhagen (VACV-Cop), given in Table S3^38^. These six non-synonymous changes are consistent with outbreak sequences in other studies^39^. Of these six genes, two have additional substitutions. MPXVgp079, an entry/fusion complex component, has an additional substitution (R194H) in 13 of our sequences, and a G4E substitution in sequence ChVir28149. MPXVgp182, a surface glycoprotein^37^, has five additional non-synonymous changes shared among the sequences from seven patients. Of the seven just-mentioned genes with non-synonymous substitutions in at least eight sequences, four (MPXVgp026, MPXVgp157, MPXVgp163, and MPXVgp182) are in genes associated with immunomodulation or host range in other orthopoxviruses^40,41^. One or two substitutions were present in 34 other genes in one to three sequences (Digital supplement, “Substitutions” sheet). No deletion of start codons or truncation of genes by the introduction of stop codons was detected.

### Gene duplication and deletion

A mapping of sequencing reads from sample ChVir28389 against another outbreak reference (ON563414.2) showed an absence of coverage in a region of ~2000 nt at the 3’ end of the genome, involving genes MPXVgp184 through MPXVgp187 (Fig. 2A). To determine the genome sequence in this region, strain-specific hemi-nested primer sets were designed to match the flanking regions so as to amplify the unknown intermediate. Primer binding sites of the forward primers were unique in the genome and excluded binding with any complementary genome region. The resulting PCR amplicons were Sanger sequenced. This revealed that a region of 856 nts from the 5’ ITR region of the genome (start site 6484) had been duplicated (reverse-complemented) to near the 3’ end of the genome (start site 188,699), replacing a region of typical length ~2048 nt (Figure 2B). The 856 nt fragment includes 502 nt before the MPXVgp005 gene and 135 nt after it. The duplication can therefore be expected to contain both the transcription leader and termination sequences and is likely a functional second copy of the gene. The elided terminal genome region results in the truncation of MPXVgp184 (20 of 106 total amino acids are still present at the 5’ end), the complete deletion of MPXVgp185 and MPXVgp186, and the truncation of MPXVgp187 (51 of 177 total amino acids remain at the 3’ end). The identical change was present in a sample from a second lesion from the same patient. We successfully isolated virus from the two lesions with the translocation. Examining pre-2022 full MPXV genome sequences, we found translocations between the same genome regions in 20 west African sequences, in a variety of host species, dating between 1958 and 2018, as had been noted for the 2003 US outbreak^2^. However, in these 20 cases the duplication direction was the opposite, from near the 3’ end of the genome to near the 5’ end, with the opposite result: duplication of MPXVgp184 through MPXVgp187 and deletion of MPXVgp005 (Fig. 2C). Although the changes in these 20 sequences are very similar, there are many small differences in affected genome positions and the length of the duplicated region (Table S4). On July 10, 2022 we downloaded^30^ 330 sequences from the 2022 outbreak and checked for either the change we see in ChVir28389 (Figure 2B) or the just-mentioned change in the other direction (Figure 2C). No other 2022 outbreak sequence had either change.

**Figure 2:**
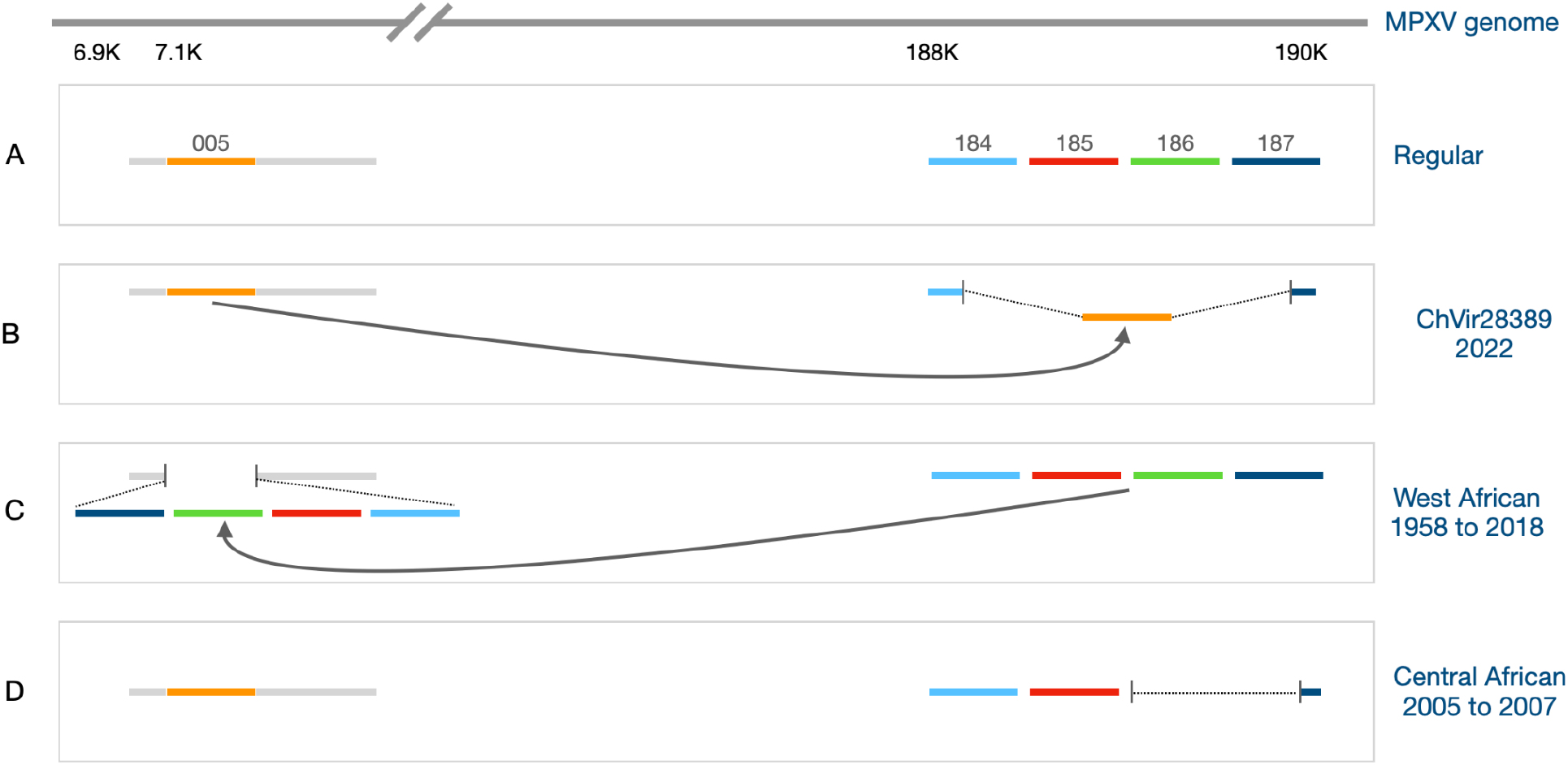
Schematic of the ChVir28389 translocation and other changes in the same genes. Genome offsets and gene lengths are not to scale. MPXV gene names are given as numbers without the “MPXVgp” prefix. Dotted regions bounded by short vertical lines indicate deletions. Arrows show the direction of duplications. **A)** The regular MPXV genome organization of MPXVgp005 and MPXVgp184-187. **B)** The translocation in ChVir28389 (and in ChVir28388, a sequence from a second lesion of the same patient). A region of 856 nt from the beginning of the genome (005, shown in orange) is duplicated (reverse complemented) at the end of the genome. The 856 nt fragment includes 502 nt before the MPXVgp005 gene and 135 nt after it and can be expected to contain both the transcription leader and termination sequences and is likely a functional second copy of the gene. The duplication resulted in the truncation of MPXVgp184 (shown in light blue, with 20 of 106 total amino acids are still present at the 5’ end), the complete deletion of MPXVgp185 (red) and MPXVgp186 (green), and the truncation of MPXVgp187 (shown in dark blue, with 51 of 177 total amino acids remaining at the 3’ end). **C)** The change observed in 20 west African sequences, as also noted for the 2003 US outbreak^2^. This change is conceptually the opposite of the change in **A**. Here, the reverse complement of the MPXVgp184-187 region is duplicated from the 3’ end of the genome to the 5’, resulting in the deletion of MPXVgp005. **D)** Deletion of MPXVgp187 and partial truncation of MPXVgp187 was found in four human MPXV cases in the Democratic Republic of the Congo in the period November 2005 to November 2007^20^.

## Discussion

Changed host population conditions, including absence of VARV- or VACV-derived immunity in those aged under ~45 years and an altered contact pattern in the involved index population, now appear to favor sustained human-to-human transmission, which is a novelty in MPXV^3,42^. There is evidence that poxvirus specialization in a new host involves inactivation of host-specific genes influencing host range, immunomodulation, and virulence^16,17^. The ongoing outbreak raises concerns regarding the potential of virus establishment in the human population. Among the most pressing issues in surveillance is the identification of potential markers of human adaptation. The dearth of genome information from humans infected prior to the present outbreak is a major challenge. The NCBI database query retrieved 76 complete pre-2022 MPXV genomes. This offers only a very modest and uneven representation of the genetic diversity of MPXV in the 64 years since its original detection in 1958. In addition to the paucity of data, a combination of factors makes it currently impossible to draw unequivocal conclusions regarding the evolution of MPXV. These include the high likelihood of multiple non-human interspecies transmissions, multiple zoonotic spillovers and possibly reverse-zoonosis, the possibility of multiple reservoir species with differing evolutionary rates, latent subclinical infections (already documented for several orthopoxviruses, possibly including MPXV^43^), and the distorting influence of genome editing by host enzymes from the APOBEC family.

It has been suggested that the MPXV genome in recent human infections may be subject to APOBEC editing during genome replication, occurring soon after zoonotic transfer from rodents, accounting for the extreme GA→AA and TC→TT frequency bias in outbreak sequence SNPs^36,37,44^. Phylogenetic anomalies in MPXV sequence similarity, root-to-tip distance, and dating could be due to the length of the post-zoonosis transmission chain at the time of sampling. A sequence from a human sample several transmission steps after a zoonosis might show a large number of nt differences as compared to one from the same date but from a human with a primary zoonotic infection. For example, in Fig. 1, many sequences from the 1970s or earlier are placed as close siblings and with similar root-to-tip distances as sequences from 2003 or later. Similarly, a sample from an early date (e.g., HM172544.1 from 1979) might have been obtained late in a transmission chain after heavy APOBEC editing, and thus be placed further from the presumed root of a phylogenetic tree than a sample obtained much later (e.g., HQ857562.1 in 2011) but immediately following zoonosis, before APOBEC editing has occurred.

Given the propensity of orthopoxviruses, including MPXV, for rapid adaptation to new hosts via duplication and alteration of terminal genes, and the unprecedented scope of opportunity for MPXV in the current situation, the duplication, inactivations, and deletions we see in five terminal genes of ChVir28389 are of great interest. The phenotypic effect, if any, of these changes cannot be inferred from the literature, most crucially because relevant functionality, when known, has usually been determined in related viruses (typically VACV), typically in mice or cell culture experiments. Further, knowledge of human MPXV has mainly been accrued for the genetically distinct central African virus‘^22^, necessarily in *post facto* investigations. Finally, the novel phenotypic characteristics of the current outbreak, such as atypical clinical presentation and sustained human-to-human transmission, directly indicate that some pre-2022 knowledge of west African MPXV may no longer apply.

The duplicated region of 856 nt in ChVir28389 contains MPXVgp005, of just 64 amino acids, typically annotated as producing a host range protein with unknown function. The annotation in the Zaire-96-I-16 full genome (NC_003310.1) and various others also adds “similar to vaccinia virus strain Copenhagen C7L” but we can find no basis for that claim. On the contrary, it has also been suggested that MPXVgp005 should not be annotated at all^24^.

Two genes are likely inactivated (MPXVgp184/187) and two are deleted (MPVXgp185/186) in ChVir28389. MPXVgp184 has been described as an apoptosis inhibitor, a transmembrane protein of the Golgi apparatus^45^. MPXVgp185 is an ortholog of VACV Copenhagen B22R/C16L (75% amino acid identity) and Cowpox (CPXV) Gri-90 D5L (93% amino acid identity). VACV C16L disrupts recognition of double-stranded RNA by DNA-PK, thereby inhibiting IRF3 activation, as demonstrated *in vivo* in mice^40,46^. CPXV D5L plays a similar role, and its mRNA transcripts have been found in abundance in virions^47^. The function of MPXVgp186 is unknown^20^. MPXVgp187 is an orthopoxvirus major histocompatibility complex class I-like protein that inhibits NKG2D-dependent killing by NK cells^48^ and whose repeated deletion in cases of human-to-human transmission in central African MPXV cases has been suggested as indicating an association with increased fitness in humans^20^.

Interestingly, changes affecting similar genome regions have been observed in other MPXV sequences. There is a counterpart in central African human sequences in which all of MPVXgp186 and the first 103 nt of MPVXgp187 (including its leader sequence) were deleted (Fig. 2D)^20^. The 20 other west African MPXV sequences in which we identified a similar change but from the 3’ end of the genome to the 5’ end (Fig. 2C), involved human and non-human host species (Table S4). Undersampling precludes an unambiguous conclusion regarding whether these changes are all independent or descend from a smaller number of unsequenced common ancestors. Recombination involving these regions may occur regularly in a stochastic fashion during genome replication, independent of selection pressure, with host-range gene variants potentially conferring a considerable spontaneous immunological advantage to the virus following spillover to a new host.

The dynamics of previous MPXV outbreaks may have no relevance to future evolution now that the virus has apparently achieved relatively sustained and widespread human-to-human transmission. The consequence of changes in poxvirus genes whose products are no longer required in a new host or otherwise altered context is unpredictable^49^. For example, genes promoting virulence in VACV are inactivated in VARV, yet VARV is much more virulent^50–52^, and the loss or inactivation of host immune system-modulating genes in VACV can result in increased virulence^40^. We should not be complacent regarding the current outbreak, based on the prior history of MPXV in humans. The phenotypic potential of a poxvirus finding itself in a new host population should not be discounted, as evidenced by the impact of the introductions of myxoma virus in Australia and squirrelpox in the UK. The poxvirus that eventually became the human-specific VARV was also originally a non-human virus. As with MPXV, VARV may have had a long early history of dead-end zoonoses; it has been convincingly argued that VARV caused only mild disease prior to the seventeenth century^53^. The MPXV phenotype we have known for the last 64 years may not resemble near-future human MPXV.

## Supporting information

Substitutions, coverage, identity metadata

## Acknowledgements

We thank Petra Mackeldanz and Patricia Tscheak for help with sample logistics. Thanks to the many laboratories worldwide who have contributed MPXV genomes.

## Ethical statement

Research clearance for the use of routine data from anonymized subjects is provided under paragraph 25 of the Berlin Landeskranken hausgesetz.

## Data sharing

Full genome sequences have been uploaded to the GISAID EpiPox database (https://www.gisaid.org/). The correspondence between ChVir identifiers used in this paper and GISAID accession ids is given in Table S5. GISAID FASTA sequence identifiers have the form hMpxV/Germany/BE-ChVirXXXXX/2022. Raw sequencing data can be obtained via request to the corresponding author.

## Supplementary Tables

**Table S1:** Sample ids of sequences generated in this study and accession numbers for sequences used in Fig. 1.

| Sample source (count) | Sample id / accession |
| --- | --- |
| Charité outbreak (48) | ChVir28136, ChVir28138, ChVir28146, ChVir28148, ChVir28149, ChVir28152, ChVir28154, ChVir28209, ChVir28220, ChVir28274, ChVir28292, ChVir28297, ChVir28305, ChVir28388, ChVir28389, ChVir28396, ChVir28423, ChVir28443, ChVir28446, ChVir28451, ChVir28452, ChVir28456, ChVir28471, ChVir28473, ChVir28474, ChVir28531, ChVir28534, ChVir28539, ChVir28566, ChVir28581, ChVir28599, ChVir28601, ChVir28604, ChVir28633, ChVir28656, ChVir28675, ChVir28679, ChVir28682, ChVir28684, ChVir28685, ChVir28689, ChVir28692, ChVir28695, ChVir28700, ChVir28703, ChVir28705, ChVir28707, ChVir28718 |
| Non-Charité outbreak (83) | MPX_37, MPX_42, ON563414.3, ON568298.1, ON585029.1, ON585030.1, ON585031.1, ON585032.1, ON585033.1, ON585034.1, ON585035.1, ON585036.1, ON585037.1, ON585038.1, ON595760.2, ON602722.2, ON609725.2, ON614676.1, ON615424.1, ON619835.2, ON619836.2, ON619837.2, ON619838.2, ON622713.1, ON622718.1, ON622720.1, ON622721.1, ON622722.2, ON627808.1, ON631963.1, ON637938.1, ON637939.1, ON644344.1, ON645312.1, ON649708.1, ON649709.1, ON649710.1, ON649711.1, ON649712.1, ON649713.1, ON649714.1, ON649715.1, ON649716.1, ON649717.1, ON649718.1, ON649719.1, ON649720.1, ON649721.1, ON649722.1, ON649723.1, ON649724.1, ON649725.1, ON649879.1, ON674051.1, ON675438.1, ON676703.1, ON676704.1, ON676705.1, ON676706.1, ON682263.2, ON682264.2, ON682265.1, ON682266.1, ON682267.1, ON682268.1, ON682269.2, ON682270.1, ON694329.1, ON694330.1, ON694331.1, ON694332.1, ON694333.1, ON694334.1, ON694335.1, ON694336.1, ON694337.1, ON694338.1, ON694339.1, ON694340.1, ON694341.1, ON694342.1, ON720848.1, ON720849.1 |
| Pre-2022 outbreak (88) | AF380138.1, AY603973.1, AY741551.1, AY753185.1, DQ011153.1, DQ011154.1, DQ011155.1, DQ011156.1, DQ011157.1, HM172544.1, HQ857562.1, JX878407.1, JX878408.1, JX878409.1, JX878410.1, JX878411.1, JX878412.1, JX878413.1, JX878414.1, JX878415.1, JX878416.1, JX878417.1, JX878418.1, JX878419.1, JX878420.1, JX878421.1, JX878422.1, JX878423.1, JX878424.1, JX878425.1, JX878426.1, JX878427.1, JX878428.1, JX878429.1, KC257460.1, KJ642613.1, KJ642615.1, KJ642616.1, KJ642617.1, KJ642619.1, KP849469.1, KP849470.1, KP849471.1, MK783028.1, MK783029.1, MK783030.1, MK783031.1, MK783032.1, MN346690.1, MN346692.1, MN346694.1, MN346695.1, MN346696.1, MN346698.1, MN346699.1, MN346700.1, MN346702.1, MN648051.1, MT903337.1, MT903338.1, MT903339.1, MT903340.1, MT903342.1, MT903343.1, MT903344.1, MT903345.1, MT903346.1, MT903347.1, MT903348.1, NC_003310.1, NC_063383.1, ON622712.1, ON631241.1, ON676707.1, ON676708.1, ON736420.1 |
| Pre-2022 outbreak omitted (7) | HQ857563.1, KC257459.1, KJ642612.1, KJ642614.1, KJ642618.1, MN346693.1, MT903341.1 |

**Table S2:** RT-PCR Cycle threshold values and sequencing statistics of 48 patient sequences. The columns give the sequence name, the RT-PCR cycle threshold (Ct) value, the total number of sequencing reads obtained (the sum of R1 and R2 counts for paired-end sequencing), the number of reads “on target” (matching MPXV outbreak sequence ON563414.2 using sensitive nucleotide matching in Geneious), and the min/max/mean coverage depth. Where the read count column has two values, the upper is for MiniSeq processing and the lower for NextSeq. Additional sequence detail is given in the Digital Supplement.

| Sequence | Ct-value (OPXV<br>TIB Molbiol) | Read count | Reads on target | min/max/mean coverage depth |
| --- | --- | --- | --- | --- |
| ChVir28136 | 23.99 | 6,677,806 | 24,691 | 1/66/18.7 |
| ChVir28138 | 18.92 | 6,800,138 | 1,427,214 | 1/4,522/1,089 |
| ChVir28146 | 20.11 | 10,819,618 | 43,742 | 1/98/32.7 |
| ChVir28148 | 18.55 | 7,040,634 | 55,934 | 1/164/42.2 |
| ChVir28149 | 24.84 | 7,251,228 | 40,490 | 1/108/30.7 |
| ChVir28152 | 24.73 | 11,051,166 | 47,178 | 1/145/35.8 |
| ChVir28154 | 20.75 | 9,552,872 | 132,082 | 1/240/99.8 |
| ChVir28209 | 23.19 | 8,719,682 | 117,924 | 1/268/88.9 |
| ChVir28220 | 17.75 | 3,375,686<br>1,761,814 | 25,576 | 0/196/16.6 |
| ChVir28274 | 20.12 | 2,567,252<br>1,136,316 | 29,212 | 1/298/18.7 |
| ChVir28292 | 19.91 | 2,429,418<br>3,579,462 | 26,940 | 0/311/15 |
| ChVir28297 | 22.00 | 3,044,394<br>5,628,078 | 12,167 | 0/309/6.7 |
| ChVir28305 | 20.81 | 6,462,850<br>5,624,418 | 845,184 | 1/8,235/658.6 |
| ChVir28388 | 24.62 | 23,564,962 | 57,422 | 1/118/43.9 |
| ChVir28389 | 19.10 | 23,566,064 | 105,983 | 0/1,978/45.8 |
| ChVir28396 | 21.32 | 35,353,730 | 263,395 | 2/709/102.5 |
| ChVir28423 | 25.67 | 17,222,870 | 77,522 | 0/725/31 |
| ChVir28443 | 22.43 | 12,510,304 | 43,632 | 0/1,203/19.6 |
| ChVir28446 | 20.99 | 11,464,154 | 106,719 | 2/2,047/45.7 |
| ChVir28451 | 25.7 | 15,863,976 | 250,323 | 0/2,284/100.4 |
| ChVir28452 | 15.82 | 1,741,360 | 12,372 | 0/448/5.3 |
| ChVir28456 | 21.92 | 7,041,554 | 32,712 | 1/447/25 |
| ChVir28471 | 27.16 | 15,791,640 | 111,463 | 0/3,594/49.7 |
| ChVir28473 | 20.16 | 27,022,776 | 169,358 | 3/4,854/80 |
| ChVir28474 | 20 | 6,550,154 | 98,414 | 3/1,183/39.9 |
| ChVir28531 | 24.78 | 22,206,746 | 12,886 | 0/444/6 |
| ChVir28534 | 21.14 | 10,284,026 | 18,485 | 0/1,072/9.1 |
| ChVir28539 | 19.89 | 10,633,496 | 26,917 | 0/1,133/13 |
| ChVir28566 | 25.77 | 3,484,626 | 90,243 | 0/825/68.9 |
| ChVir28581 | 27.91 | 30,587,788 | 57,729 | 2/433/43.6 |
| ChVir28599 | 16.25 | 3,659,858 | 37,896 | 1/205/28.8 |
| ChVir28601 | 16.69 | 2,106,808 | 12,258 | 0/55/9.3 |
| ChVir28604 | 22.45 | 4,304,994 | 58,317 | 1/833/44.6 |
| ChVir28633 | 20.31 | 24,272,690 | 57,483 | 1/127/43.7 |
| ChVir28656 | 21.09 | 18,462,182 | 10,335 | 0/30/7.8 |
| ChVir28675 | 26.79 | 21,088,062 | 17,080 | 0/43/13 |
| ChVir28679 | 18.69 | 11,003,042 | 26,857 | 1/60/19.8 |
| ChVir28682 | 27.63 | 22,138,092 | 108,856 | 1/249/82.9 |
| ChVir28684 | 19.88 | 33,669,658 | 118,241 | 1/217/89.4 |
| ChVir28685 | 23.06 | 24,175,248 | 54,552 | 1/133/41.4 |
| ChVir28689 | 18.68 | 31,239,686 | 515,383 | 1/1,010/391.7 |
| ChVir28692 | 24.03 | 8,099,774 | 17,819 | 0/276/13.6 |
| ChVir28695 | 21.94 | 6,605,766 | 11,271 | 0/167/8.6 |
| ChVir28700 | 21.55 | 9,054,896 | 11,057 | 0/85/8.4 |
| ChVir28703 | 24.7 | 7,226,170 | 39,498 | 1/259/30.1 |
| ChVir28705 | 16.77 | 18,622,244 | 70,015 | 1/505/53.4 |
| ChVir28707 | 21.44 | 8,570,860 | 53,890 | 1/724/41.1 |
| ChVir28718 | 17.92 | 4,594,098 | 35,791 | 1/137/27.3 |

**Table S3:** MPXV virus homologous gene names and notes. The GenBank ‘note’ field from the MPXV ON676708.1 record is given in the middle column, corresponding to the numeric MPXV gene name (left column). Additional notes are in the right column.

| MPXV name | ON676708.1 GenBank record 'note' field | Other notes |
| --- | --- | --- |
| MPXVgp005 | D2L; Host range protein | Appears to be incorrectly annotated in Zaire-96-I-16 (NC_003310.1) as "similar to VACV Copenhagen C7L". |
| MPXVgp045 | Palmitylated EEV membrane glycoprotein (Cop-F13L) C19L phospholipase D-like similar to Vaccinia virus strain Copenhagen F13L major envelope antigen of EEV wrapping of IMV to form IEV |  |
| MPXVgp078 | G9R VLTf-1 (late transcription factor 1) (Cop-G8R) late gene transcription factor VLTf-1 similar to Vaccinia virus strain Copenhagen G8R |  |
| MPXVgp079 | Entry/fusion complex component myristylprotein (Cop-G9R) G10R myristylated protein similar to Vaccinia virus strain Copenhagen G9R |  |
| MPXVgp083 | M4R similar to Vaccinia virus strain Copenhagen L4R ss/dsDNA binding protein (VP8) (Cop-L4R) ssDNA binding stimulation of I8R helicase activity virion core protein |  |
| MPXVgp157 | A47R IL-1/TLR signaling inhibitor (Cop-A46R) similar to Vaccinia virus strain Copenhagen A46R |  |
| MPXVgp182 | B21R Surface glycoprotein cadherin-like domain putative membrane-associated glycoprotein |  |
| MPXVgp184 | R1R; NMDA receptor-like protein | Not clear what R1R refers to, since there is no R1R in VACV Copenhagen. Referred to as an "Apoptosis inhibitor, transmembrane protein of Golgi apparatus" in ref<sup>45</sup>. |
| MPXVgp185 | Hypothetical protein (Cop-C16L) N1R VAC B15R-like similar to Vaccinia virus strain Copenhagen B22R |  |
| MPXVgp186 | MPV-Z-N2R |  |
| MPXVgp187 | N3R; MPV-Z-N3R | OMCP in ref <sup>48</sup>. |

**Table S4:** Gene duplication and deletion in regions similar to those in ChVir28389. A similar pattern of gene duplication to that seen in ChVir28389 is present in 20 MPXV sequences from the period 1958-2018. The first four columns give the GenBank accession number, country abbreviation, sampling year, and host species. Then three columns indicate where the duplicated region is copied to (with the start and stop nucleotide locations and the length of the duplication). The final three columns indicate the source of the copy. Note that in the case of KJ642616.1, the region that is duplicated (2297 nt) is three nucleotides shorter than the region present at the start of the genome (2300 nt). This pattern of duplication is not identical to that seen in ChVir28389 because the direction of the duplication in the above table is the opposite (from the genome end to its beginning) and these duplications (of genes MPXVgp184 through MPXVgp187) result in the deletion of MPXVgp005 (Fig. 2C) whereas in ChVir28389, MPXVgp005 is duplicated at the end of the genome, likely inactivating MPXVgp184 and MPXVgp187 and deleting MPXVgp185 and MPXVgp186 (Fig. 2B). Species abbreviations are CH: Chimpanzee, CM: Cynomolgus monkey, DM: Dormouse, HS: Homo sapiens, PD: Prairie dog, PR: Gambian pouched rat, RS: Rope squirrel.

| Accession | Country | Year | Host species | To |  |  | From |  |  |
| --- | --- | --- | --- | --- | --- | --- | --- | --- | --- |
|  |  |  |  | Start | Stop | Length | Start | Stop | Length |
| AY753185.1 | DK | 1958 | CM | 6554 | 8850 | 2297 | 190620 | 192916 | 2297 |
| AY603973.1 | WA | 1961 | CM | 6452 | 8748 | 2297 | 190448 | 192744 | 2297 |
| KJ642616.1 | FR | 1968 | HS | 6266 | 8565 | 2300 | 190180 | 192476 | 2297 |
| DQ011156.1 | LR | 1970 | HS | 7063 | 9356 | 2294 | 190901 | 193194 | 2294 |
| AY741551.1 | SL | 1970 | HS | 6276 | 8572 | 2297 | 190185 | 192481 | 2297 |
| KP849470.1 | CI | 1971 | HS | 7232 | 9525 | 2294 | 190943 | 193236 | 2294 |
| DQ011153.1 | US | 2003 | PD | 6538 | 8835 | 2298 | 189946 | 192243 | 2298 |
| DQ011157.1 | US | 2003 | HS | 6538 | 8835 | 2298 | 189946 | 192243 | 2298 |
| MT903346.1 | US | 2003 | PR | 6537 | 8834 | 2298 | 189945 | 192242 | 2298 |
| MT903347.1 | US | 2003 | DM | 6538 | 8835 | 2298 | 189946 | 192243 | 2298 |
| MT903348.1 | US | 2003 | RS | 6538 | 8835 | 2298 | 189946 | 192243 | 2298 |
| MN346690.1 | CI | 2017 | CH | 6213 | 8504 | 2292 | 189947 | 192238 | 2292 |
| MN346692.1 | CI | 2017 | CH | 6213 | 8504 | 2292 | 189918 | 192209 | 2292 |
| MN346694.1 | CI | 2017 | CH | 6213 | 8504 | 2292 | 189912 | 192203 | 2292 |
| MN346695.1 | CI | 2017 | CH | 6213 | 8504 | 2292 | 189867 | 192158 | 2292 |
| MN346696.1 | CI | 2017 | CH | 6213 | 8504 | 2292 | 189871 | 192162 | 2292 |
| MN346698.1 | CI | 2017 | CH | 6213 | 8504 | 2292 | 189804 | 192095 | 2292 |
| MN346699.1 | CI | 2017 | CH | 6213 | 8504 | 2292 | 189804 | 192095 | 2292 |
| MN346700.1 | CI | 2017 | CH | 6213 | 8504 | 2292 | 189804 | 192095 | 2292 |
| MN346702.1 | CI | 2018 | CH | 6199 | 8490 | 2292 | 189816 | 192107 | 2292 |

**Table S5:** Charité sequence identifiers and corresponding GISAID accession.

| **Charité id** | **GISAID accession** |
| --- | --- |
| ChVir28136 | EPI_ISL_13889442 |
| ChVir28138 | EPI_ISL_13890273 |
| ChVir28146 | EPI_ISL_13890481 |
| ChVir28148 | EPI_ISL_13890467 |
| ChVir28149 | EPI_ISL_13889446 |
| ChVir28152 | EPI_ISL_13889660 |
| ChVir28154 | EPI_ISL_13889436 |
| ChVir28209 | EPI_ISL_13890474 |
| ChVir28220 | EPI_ISL_13889447 |
| ChVir28274 | EPI_ISL_13890475 |
| ChVir28292 | EPI_ISL_13890135 |
| ChVir28297 | EPI_ISL_13889729 |
| ChVir28305 | EPI_ISL_13889908 |
| ChVir28388 | EPI_ISL_13890338 |
| ChVir28389 | EPI_ISL_13890482 |
| ChVir28396 | EPI_ISL_13890476 |
| ChVir28423 | EPI_ISL_13889515 |
| ChVir28443 | EPI_ISL_13890464 |
| ChVir28446 | EPI_ISL_13890471 |
| ChVir28451 | EPI_ISL_13889796 |
| ChVir28452 | EPI_ISL_13889437 |
| ChVir28456 | EPI_ISL_13889977 |
| ChVir28471 | EPI_ISL_13890480 |
| ChVir28473 | EPI_ISL_13889445 |
| ChVir28474 | EPI_ISL_13889441 |
| ChVir28531 | EPI_ISL_13890470 |
| ChVir28534 | EPI_ISL_13890472 |
| ChVir28539 | EPI_ISL_13889450 |
| ChVir28566 | EPI_ISL_13890048 |
| ChVir28581 | EPI_ISL_13889448 |
| ChVir28599 | EPI_ISL_13890468 |
| ChVir28601 | EPI_ISL_13890479 |
| ChVir28604 | EPI_ISL_13889590 |
| ChVir28633 | EPI_ISL_13890465 |
| ChVir28656 | EPI_ISL_13889435 |
| ChVir28675 | EPI_ISL_13889443 |
| ChVir28679 | EPI_ISL_13889438 |
| ChVir28682 | EPI_ISL_13890469 |
| ChVir28684 | EPI_ISL_13889449 |
| ChVir28685 | EPI_ISL_13890473 |
| ChVir28689 | EPI_ISL_13889439 |
| ChVir28692 | EPI_ISL_13890204 |
| ChVir28695 | EPI_ISL_13890477 |
| ChVir28700 | EPI_ISL_13890466 |
| ChVir28703 | EPI_ISL_13890408 |
| ChVir28705 | EPI_ISL_13889444 |
| ChVir28707 | EPI_ISL_13889440 |
| ChVir28718 | EPI_ISL_13890478 |

## Supplementary digital data

Supplementary digital data is provided as an Excel spreadsheet with sheets, as follows.

### ‘Substitutions’: Non-synonymous changes in outbreak sequences

Of the sequences presented here, 38 had genome coverage of at least 99.95% and were closely examined for non-synonymous nucleotide substitutions in translated regions of the genome. The columns give: **A**) the MPXV gene name, the start (**B**) and stop (**C**) locations of the gene (from the GenBank annotation of ON676708.1), **D**) the set of substitutions, **E**) the number of sequences that have exactly that set of substitutions, **F**) the names of the sequences that all have that set of substitutions (with the “ChVir28” prefix removed for conciseness), and **G**) the note field from the ON676708.1 annotation. Note a substitution may appear in several sets for the same gene. For example, E353K is listed twice for gene MPXVgp045 because it occurs once with D294N in sequence ChVir28675 and occurs in isolation in the 37 other high-coverage sequences.

### ‘Coverage’: Genome coverage, GC content, and nucleotide composition

Columns give (**A**) the name of the sequence, (**B**) the sequence length in nucleotides, (**C**) the number of genome locations with no coverage, (**D**) the coverage fraction, (**E**) the GC nucleotide fraction, the genome count of individual nucleotides (columns **F** through **I**: ACGT), missing coverage (column **J**, marked ?) according to Geneious (version 2022.0.1), and ambiguous nucleotides (columns **K** through **R**).

### ‘Identity’: Full genome pairwise-aligned sequence nucleotide (nt) identity

The leftmost column shows the name of outbreak sequences from Berlin. The numbers in the cells in the top row correspond to numbered sequences in the first column, plus the nearest pre-outbreak sequence (MT903344.1) in the rightmost column. The nt identity fraction is computed using a numerator that is the sum of the number of identical nts and the number of ambiguously-matching nts, and a denominator that is the length of the shorter of the two sequences, minus the number of non-covered sites.

### ‘Identity detail’: Detailed full genome pairwise-aligned sequence nucleotide (nt) identity

The identity is computed as in the ‘Identity’ sheet (described above). Other cell detail is according to the following abbreviations L: sequence Length; G: number of Gaps in sequence; C: number of no Coverage characters in sequence; N: number of fully-ambiguous N characters in sequence; IM: Identical nt Matches; AM: Ambiguous nt Matches; GG: Gap/Gap matches (both sequences have gaps); G?: Gap/Non-gap mismatches (one sequence has a gap); CC: No coverage/No coverage (both sequences have no coverage); C?: No coverage (one sequence has no coverage); NE: Non-equal nt mismatches. Note that the sequence information in the left-most column is for the unaligned sequence, whereas that in the other cells describes the alignment between two sequences. For example, neither ChVir28388 nor ChVir28389 have any gap characters in their sequences (so “G:0” appears in the left column), in the pairwise alignment there are 142 gaps (and “G:142” in the cell at their intersection in the table).

### ‘GISAID accession’: Correspondence between Charité ids and GISAID accession ids

As also given in Table S5.

